# Single cell genome sequencing of laboratory mouse microbiota improves taxonomic and functional resolution of this model microbial community

**DOI:** 10.1101/2021.12.13.472402

**Authors:** Svetlana Lyalina, Ramunas Stepanauskas, Frank Wu, Shomyseh Sanjabi, Katherine S. Pollard

## Abstract

Laboratory mice are widely studied as models of mammalian biology, including the microbiota. However, much of the taxonomic and functional diversity of the mouse gut microbiome is missed in current metagenomic studies, because genome databases have not achieved a balanced representation of the diverse members of this ecosystem. Towards solving this problem, we used flow cytometry and low-coverage sequencing to capture the genomes of 764 single cells from the stool of three laboratory mice. From these, we generated 298 high-coverage microbial genome assemblies, which we annotated for open reading frames and phylogenetic placement. These genomes increase the gene catalog and phylogenetic breadth of the mouse microbiota, adding 135 novel species with the greatest increase in diversity to the *Muribaculaceae* and *Bacteroidaceae* families. This new diversity also improves the read mapping rate, taxonomic classifier performance, and gene detection rate of mouse stool metagenomes. The novel microbial functions revealed through our single-cell genomes highlight previously invisible pathways that may be important for life in the murine gastrointestinal tract.

## Introduction

The number of microbial species with at least one genome sequence has grown rapidly in recent years. The human gut has been a major focus of these efforts[1–5], with metagenome assembled genomes (MAGs) and innovations in culturing[6–8] capturing genomes for many species previously absent from databases built primarily through isolate sequencing.

Mice are a model system for host-associated microbiota. They are heavily utilized in biomedical research as well as basic science investigations of community assembly and resilience. However, the species present in wild and laboratory mouse stool are heavily under-represented in genome databases in comparison to human-associated microbiota[9]. This gap can create a biased picture of the functional and taxonomic landscape of shotgun metagenomic studies carried out in mice, since most bioinformatics methods rely on available reference data. Several research groups have actively sought to address this problem, both by focusing on mouse-specific bacterial strains that were previously unculturable[10] and by performing co-assembly of large-scale metagenomic datasets from a broad variety of mouse facilities[11].

This study aims to increase the number of mouse gut species with a sequenced genome using microbial single-cell genomics (SCG). Our workflow leverages fluorescence-activated cell sorting (FACS), whole genome amplification with WGA-X, shotgun sequencing and *de novo* assembly of genomes from individual microbial cells from two laboratory mouse strains[12]. By annotating the taxonomy and encoded functions of 298 quality-controlled, single-cell genomes, we revealed previously invisible pathways and phylogenetic breadth, increasing the power of metagenomic analysis tools. These results demonstrate the utility of SCG for characterizing host-associated microbiomes and provide a resource towards a better understanding of the mouse gut as a model system.

## Results

The biological material used for this study came from fecal pellets of three mice of two different strains - two wild-type C57BL/6N mice and a transgenic CD4-dnTβRII (DNR) mouse prone to developing intestinal inflammation[13]. These two strains’ intestinal microbiota have been previously studied within the lab[14], which allowed us to evaluate how the single-cell genomes we produced change previous interpretations of shotgun metagenomic data.

Using stool from these mice, we performed FACS followed by whole genome amplification with WGA-X. Cell sorting was based on the fluorescence of nucleic acids stain SYTO-9 (Thermo Fisher Scientific) and light scatter signals using a previously established gate for individual prokaryotic cells[12]. To assess the general structure of the microbiomes, we first performed low-coverage sequencing and assembly of 738 cells (median 765,918 reads/sample [342,424 - 2,670,861]) (Methods). We filtered the resulting single-cell amplified genomes (SAGs) to exclude assemblies with total length below 20,000 basepairs (bp) or suspected to be contaminated (determined by nucleotide tetramer principal components analysis[15]), producing 697 SAGs that vary in quality and completeness (Fig 1). Compared to the earlier, multiple displacement amplification (MDA) technique[16], the WGA-X approach has been shown to improve the amplification of single-cell DNA, especially for microorganisms with high GC-content genomes[12], and we indeed observed a wide range of GC% across the assemblies (Fig 1E).

**Fig 1.**
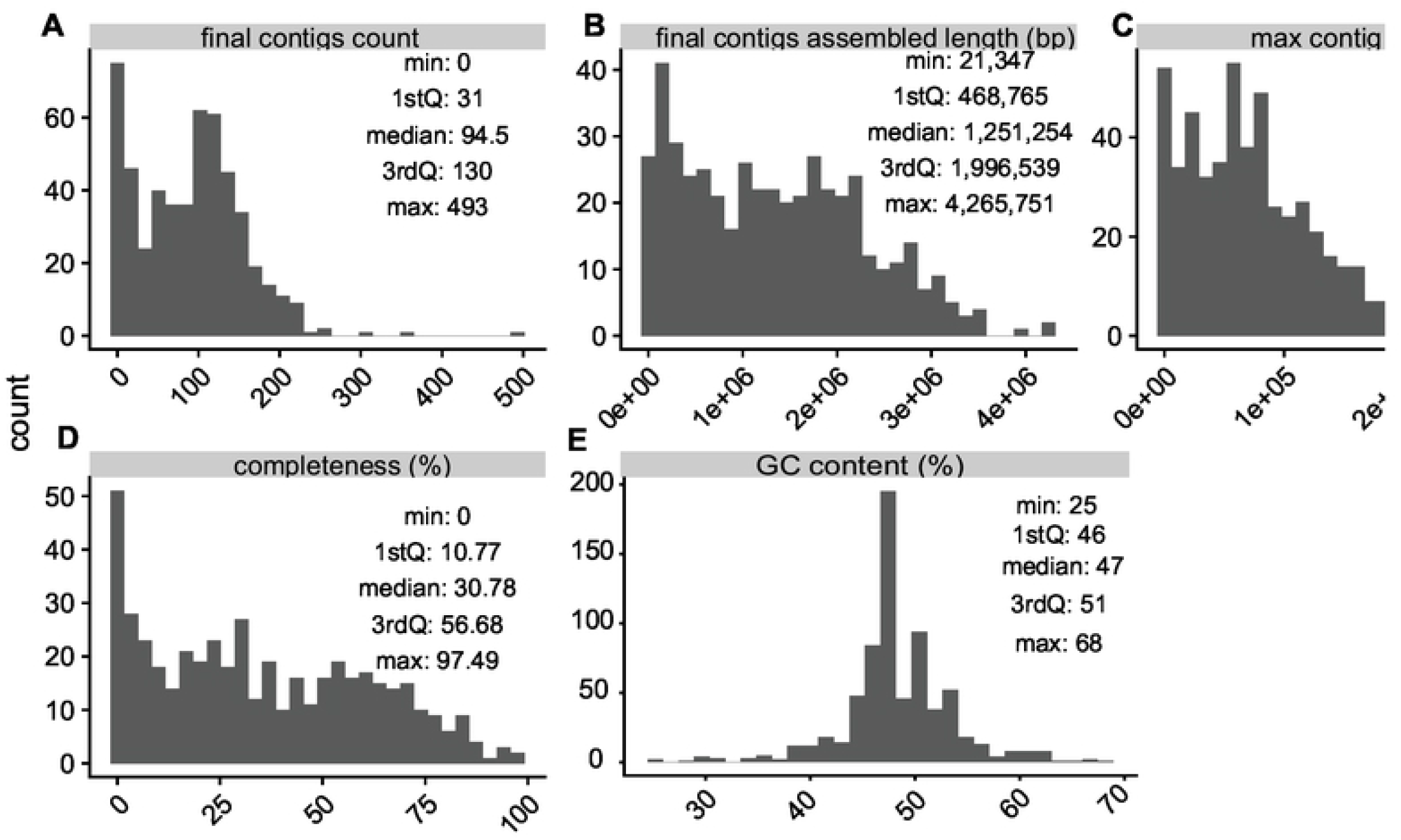
Quality metrics of low-coverage SAG assemblies. A faceted plot containing histograms of quality metrics used to describe the assembled SAGs. The facets display the following metrics: A) total number of contigs, B) their total assembled lengths (in number of nucleotide basepairs), C) the length of the longest contig in each assembly (in number of nucleotide basepairs), D) CheckM estimated completeness (as percentage), and E) GC content. Tukey five-number summaries (minimum, 25% quantile, median, 75% quantile, maximum) are overlaid on each metric’s panel.

We next selected two samples, one of each strain, for further sequencing towards obtaining high-coverage SAGs. To prioritize cells that would produce high-quality data and increase the taxonomic diversity of mouse gut genomes, we performed phylogenetic placement of the low-coverage SAGs with GTDB-Tk[17], successfully placing 448 SAGs within the GTDB genome tree of life[18] (release 86). We then selected the 150 SAGs from each sample that maximize phylogenetic diversity and excluded SAGs with low probability of high genome recovery (Methods). Further sequencing and assembly of DNA from the corresponding cells produced 298 high-coverage SAGs after quality control. As expected, these show significant improvements in relevant quality metrics when compared to corresponding low-coverage assemblies (S1 Fig). All subsequent analyses use the high-coverage SAGs.

To evaluate whether the SAGs increased the diversity of sequenced mouse microbiota, we placed them on the GTDB tree and quantified the additional branch length added by SAGs compared to the total branch length from previously sequenced microbial samples. Evaluating this metric across clades, we observed that our SAGs primarily increase the phylogenetic diversity of the *Muribaculaceae* and *Bacteroidaceae* families (Fig 2). Despite the fact that GTDB includes MAGs from uncultured microbes, this study adds substantial new diversity to the tree, with 135 out of 298 SAGs having no hit in the GTDB with FastANI similarity above 97%.

**Fig 2.**
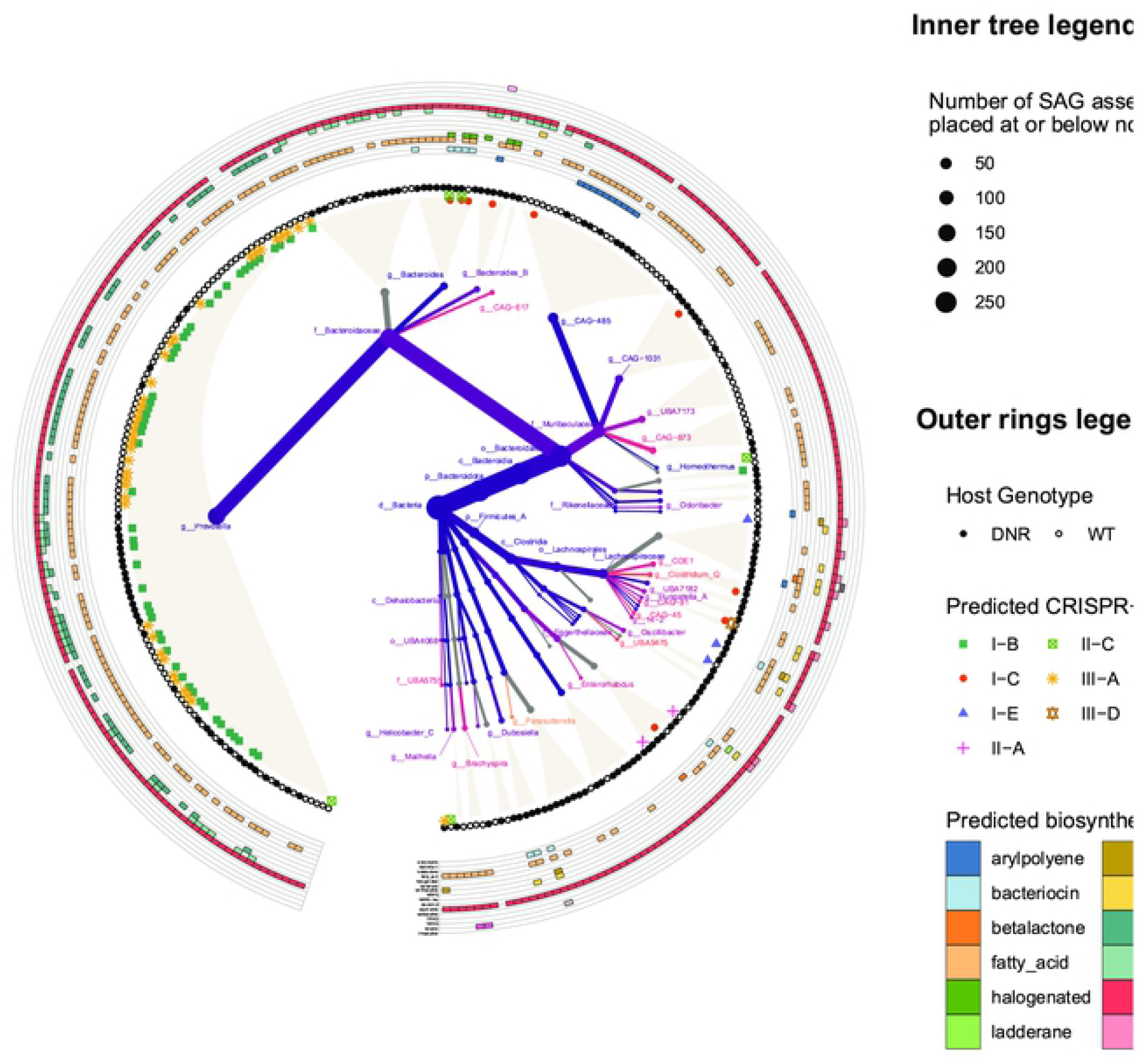
SAGs increase phylogenetic diversity and contain distinct genomic features. The central part of this circular figure contains a heat tree reflecting the number of SAG assemblies placed at different sub-branches of the GTDB v86 bacterial genome tree (represented by node size), and percentage phylogenetic gain achieved by the insertion of the new genome assemblies (represented by color scale). The outer rings of the figure contain additional genomic feature information inferred about the successfully placed SAG assemblies. The additional markings denote predicted CRISPR-Cas system type (ring of single point symbols) and the number of genes contributing to predicted biosynthetic gene clusters (outermost ring of colored polygons).

Next, we investigated the gene content of the SAGs. We annotated open reading frames in all SAGs, dereplicated these, and analyzed their functional potential using annotations from clusters of orthologous groups (COGs)[19]. Gene sequences were evaluated for percent nucleotide identity to all sequences in a previously published mouse stool metagenome-derived gene catalog (4) and labeled as novel if they have no matches above 95% nucleotide identity. Overall, 53.7% of SAG genes were novel and 46.3% overlapped with the mouse catalog, which compares to 10% overlap with a human gene catalog and <0.1% for a marine catalog (Fig 3), highlighting the functional differences of microbes across these environments. Novel SAG genes were enriched for COG categories M (Cell wall/membrane/envelope biogenesis), L (Replication, recombination and repair), C (Energy production and conversion) and R (General function prediction only). This enrichment was determined by Annotation Enrichment Analysis[20], a method that aims to reduce the bias towards highly annotated functional categories and utilize the hierarchical structure in a given functional ontology. While these annotation categories provide a rather broad summary of the functions distinct to this gene set, they generally suggest that sequencing more members of the microbiota would expand our understanding of both internal housekeeping functions (categories L and R), but also functions more pertinent for translational applications within category M, which contains potential candidates for studying interactions with the host immune system. Thus, our SAG gene catalog expands the representation of putative functions present in mouse gut microbes, with surprisingly large gains given the number of genomes sequenced for this study.

**Fig 3.**
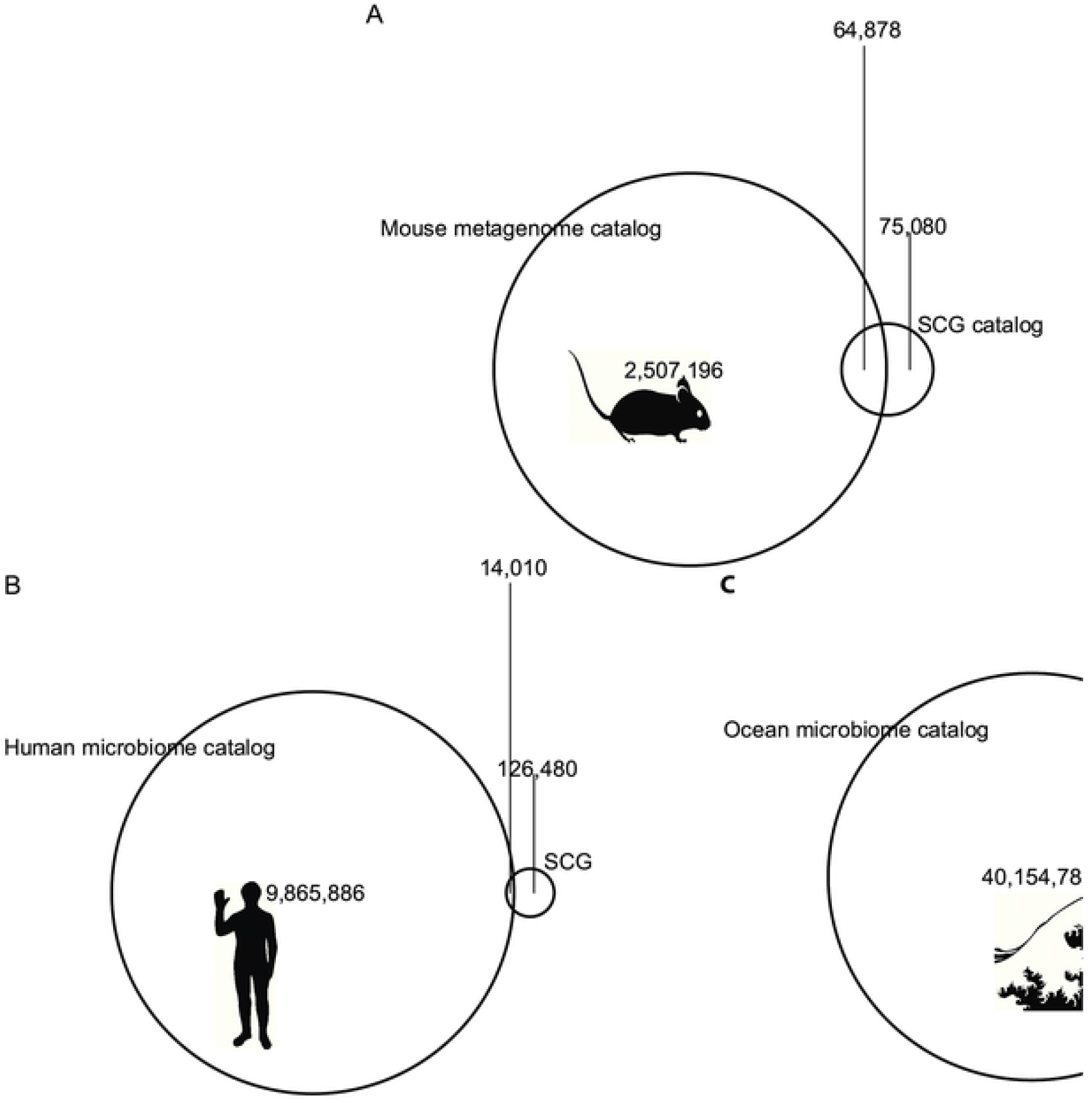
A gene catalog derived from SAGs shows subsantial novelty when compared against other microbiome gene catalogs. Euler plots reflect the shared and unique counts of genes when comparing the set of non-redundant genes from this study’s data against previously published gene catalogs derived from metagenomic sequencing efforts in A) mice, B) humans, and C) marine samples.

To expand beyond COG annotations for two important groups of genes, we performed additional annotation of enzymes involved in secondary metabolism and CRISPR associated (Cas) proteins along with their CRISPR arrays. Overall, 3,257 putative secondary metabolism gene clusters were found across the 298 SAGs sequenced at high coverage. The most prevalent predicted cluster types were the broad categories of saccharide, fatty acid, and NRPS-like, whereas the more nuanced product types were detected much more rarely. CRISPR-cas types were determined in 88 genomes, of which 22 genomes had 2 CRISPR complexes. An additional 28 genomes had Cas operons, but no proximal CRISPR array.

The distributions of biosynthetic gene clusters (BGCs) and CRISPR-Cas systems in our SAGs support the phylogenetic novelty of several clades characterized in this study. We quantified the presence of BGCs and CRISPR-cas types in relation to the phylogenetic placement of the contributing genome (outer ring of Fig 3). In this trimmed genome subtree, the newly sequenced *Prevotella* SAGs form a distinct, relatively flat phylogenetic subcluster, distinguished by unique CRISPR-Cas subtype patterns and presence of NRPS-like predicted BGCs. A closely related subset of SAGs assigned to the genus CAG-486 within the *Muribaculaceae* family accounts for a high proportion of identified aryl polyene BGCs, suggesting similar adaptations to oxidative stress[21]. Thus, the new taxonomic diversity we captured is mirrored by gene functional profiles that differ from related genomes.

Finally, we investigated to what degree our SAGs improve the sensitivity and resolution of metagenomic analysis using 236 shotgun metagenome samples from laboratory mouse stool, as well as metagenomes from wild mouse stool (N=10), human stool (N=274), and marine environments (N=20, subset of full data) (accessions listed in S2 Table). Focusing on taxonomic classifiers, we created custom mapping references for sourmash[22] and MIDAS[9], which represent two common approaches: kmer-based versus marker gene-based. We compared taxonomic coverage and prevalence estimates with each tool using the database distributed with the software, a database composed only of SAGs, and the two combined. For both tools, the combined database generally improved the taxonomic classification of mouse microbiome samples, with the exception of the wild mouse microbiome testing scenario, which only showed improvement with FDR < 0.1 when using sourmash and not MIDAS. Interestingly the addition of SAGs also improved classification rates to a limited degree with human microbiome samples (not statistically significant), but not marine samples. The results of non-parametric testing of the performance of pairs of databases for each dataset and tool type can be found in S3 Table, with highlighted rows showing cases of significant performance improvement in a number of murine shotgun microbiome datasets. Ridgeline plots graphically portray these performance differences in greater detail (S4 Fig, S5 Fig). These results show that the novel phylogenetic diversity we captured with SAGs has a positive effect on our ability to taxonomically profile shotgun metagenomes from the mammalian gut.

## Discussion

To our knowledge, this study is the first to generate single-cell genome assemblies from mammal-associated microbiota with the WGA-X approach. The draft genomes that we assembled increase the phylogenetic diversity of mouse gut microbiota in public databases. Our SAGs add a particularly large number of genomes (58 assemblies) to the recently proposed candidate family *Muribaculaceae* within the Bacteroidales, previously referred to in the literature as S24-7 and Ca. Homeothermaceae[23][24]. This family has been reported as a taxon of interest in multiple studies[25–27] but has so far only been characterized via 16S markers and MAGs. Only one recent paper has successfully isolated members of this family in culture[24]. Another taxon with large numbers of newly placed SAGs (120 assemblies), though small phylogenetic gain (4.27%), is the genus *Prevotella,* which contains Gram-negative obligate anaerobes with potential links to mucosal inflammation susceptibility[28]. Hence, our SAGs add genomes for important taxonomic groups in the mouse microbiota.

SAGs also increase our knowledge of the functional potential of microbes in the mouse gut. Gain in functional novelty includes a large number of COGs that were enriched and depleted compared to open reading frames previously observed in mouse stool samples. When summarising these differentially detected functional categories, four are particularly enriched: energy production and conversion (C), replication and repair (L), cell wall/membrane/envelope biogenesis (M), and the unspecific category (R) - general function prediction only. Previously unobserved sequences classified under the M category could be of interest when mining for new antigenic proteins, whereas genes placed in the unspecific R category could be further experimentally probed to shed light on microbial “dark matter”.

Our annotations of SAGs for secondary metabolism genes and CRISPR systems aim to highlight the capacity of this sequencing approach to more faithfully reflect intra-genome structure. When analyzed in the context of phylogenetic relationships between SAGs, the results of CRISPR-Cas type identification show SAGs placed in the *Prevotella* genus have both Type I and Type III systems, whereas this is relatively uncommon in our data outside this clade. This suggests that these microbes have a more sophisticated defense repertoire that allows for targeting of both DNA and RNA[30].

Looking at secondary metabolism, we see that the most widely represented gene clusters are for saccharide and fatty acid biosynthesis. The remaining categories are sparsely observed. An interesting clustering occurs for the resorcinol group which appears primarily to be present in genomes from the *Bacteroidaceae* family. This cluster type originates mainly from genomes found in the DNR mouse microbiome (34 resorcinol clusters predicted, vs only 6 from WT). The particular gene that is considered by the predictive tool AntiSMASH as a signature gene for the resorcinol annotation is DarB (KEGG orthology ID of K00648), which falls under the fatty acid biosynthesis KEGG pathway. The literature provides limited insight into what microbiome activities resorcinol biosynthesis could be relevant to, however, some reported associations of the more specific chemical family of dialkylresorcinols include anti-inflammatory, anti-proliferative, and antibiotic activities[31]. Interestingly, a dialkylresorcinol compound has been used to attenuate the effects of experimentally induced intestinal inflammation[32], which has potential implications for the observed higher prevalence of dialkylresorcinol-producing genomes in the inflammation-prone DNR mouse strain.

Considering the relatively modest costs of this sequencing experiment, we were surprised to find that the new sequences significantly helped with metagenomic read recruitment even in unrelated mouse lines and wild mouse samples, which have been shown to have more diverse microbiomes than their laboratory counterparts[33]. This corroborates prior reports demonstrating the value of SAG genomes as reference material for the interpretation of marine[34,35] and soil[12,36] microbiome omics data. The lack of improvement of the taxonomic classifiers on marine metagenomic data with mouse microbiome SAGs agree with our findings of novel genes, confirming the lack of highly similar genomes between these two environments.

Despite single-cell sequencing being a promising approach for increasing the representation of unculturable mouse symbionts in the tree of life, certain caveats still exist. For example, although the individual SAG assemblies have acceptable quality metrics, there is a limit to the completeness that can be achieved when operating with short read sequencing data. Long repetitive segments continue to pose an obstacle to assemblers that attempt to span ambiguous regions of the genome. Whole genome amplification, while drastically improved by the WGA-X process, is still not uniform across the genome, thus requiring a relatively deep sequencing of SAGs in order to access under-amplified regions. Despite these limitations, we expect that the taxonomic and functional novelty revealed in this study will encourage others to leverage single-cell genomics technologies.

## Materials and Methods

### Sample acquisition and sequencing

Cells were sequenced from three murine fecal pellets, two from wild-type C57BL/6N mice and one from an inflammatory bowel disease model CD4-dnTGFBRII (DNR) [13,37] mouse not exhibiting intestinal pathology at the time of sampling. To preserve the mouse feces, a cryopreservation “glyTE” stock (11.11x) was made by mixing 20 mL of 100x Tris-EDTA pH 8.0 (Sigma) with 60 mL deionized water and 100 mL molecular-grade glycerol (Acros Organics). This mixture was filter-sterilized using a 0.2 micrometer filter. Prior to use, 1x glyTE was made by diluting with phosphate buffered saline (PBS) at a 10:1 ratio. 1 mL of the 1x glyTE was then aliquoted into cryotubes. Each fecal pellet was distributed into 3 separate cryotubes to create 3 replicates for each sample. Each sample was dispersed into the solution by gentle pipetting and allowed to incubate at room temperature for 1 minute before being placed on dry ice. Samples were stored at −80 C and shipped on dry ice to the Bigelow Laboratory’s Single Cell Genomics Center for further processing using a previously described protocol[12]. Low-coverage SAG assemblies were generated to evaluate microbiome composition. Two samples, one of each murine host genotype, were selected for high-coverage sequencing. In each sample, cells were prioritized by optimizing for robust amplification profiles and maximizing the phylogenetic diversity (python code DOI: 10.5281/zenodo.2749707). The criterion used to assess amplification dynamics was computed as the time needed to reach the inflection point in the amplification curve. Raw reads were processed into assembled contigs (same procedure as described in [12]), which were further filtered to yield sufficient quality SAGs, which were assessed by checkM[38] for contamination and assigned a putative taxonomic lineage. Versions of QC and assembly pipeline subcomponents were as follows: SPAdes v3.9.0[39], bcl2fastq v2.17.1.14 (Illumina), Trimmomatic v0.32[40], kmernorm 1.05 (https://sourceforge.net/projects/kmernorm/). This SAG generation, sequencing and assembly workflow was previously evaluated for assembly errors using three bacterial benchmark cultures with diverse genome complexity and GC content (%), indicating no non-target and undefined bases in the assemblies and average frequencies of mis-assemblies, indels and mismatches per 100 kbp being 0.9, 1.8, and 4.7[12].

All mice were housed and bred in specific pathogen-fee conditions in the Gladstone animal facility. No animals were euthanized for the purposes of this study. All animal experiments were conducted with all relevant ethical regulations for animal testing and research and were done in accordance with guidelines set by the Institutional Animal Care and Use Committee of the University of California, San Francisco under protocol #AN151865–03A.

### Computational analyses of phylogenetic placement and predicted gene function

We used pplacer[41] within GTDB-Tk[17] to phylogenetically place the SAGs in the genome tree that is part of GTDB release 86. The resulting placements were used to calculate phylogenetic diversity and phylogenetic gain from the SAGs using GenomeTreeTk[42]. The heat tree visualization was inspired by the approach illustrated in the metacoder[43] R package and was ultimately generated alongside additional genomic feature annotation via the ggtree[44] and ggtreeExtra[45] packages.

Classification of the CRISPR-Cas system types and subtypes was done by CRISPRCasTyper v1.2.1[46]. Identification of secondary metabolism gene clusters was performed with AntiSMASH v5.2[47]. Unless otherwise stated, default settings were used when invoking these computational tools.

Clustering of predicted genes was performed by CD-HIT-EST v4.6.8 [48] (settings: −r 1 −c 0.95 −n 8), and the resulting gene catalog was compared by CD-HIT-EST-2D to previously published gene catalogs derived from mouse[11], human[49], and marine[50] microbiomes. To gauge enrichment of functional categories for novel sequences in our catalog, we annotated the sequences with EggNOG-mapper v1.0.3 [51] using diamond[52] as the homology search method and then applied Annotation Enrichment Analysis methodology[20] to assess the relationship between the number of genes assigned to a COG category and their novelty in relation to the previously published mouse metagenome catalog[11]. We corrected for multiple testing using the p.adjust function in base R[53] (v3.6.0), using the Benjamini-Hochberg[54] method.

### Comparative analyses of metagenomic read recruitment

Custom sourmash[22] lowest common ancestor (LCA) databases for the set of GTDB genomes and SAG assemblies were created using the “sourmash lca index” function, and metagenomic datasets were then classified with “sourmash lca summarize” using the two databases separately as well as together to evaluate the effect of combining the data. To create the relevant databases for MIDAS, we used the built-in database creation script within the package, as well as an auxiliary step of assigning certain SAG assemblies to pre-existing genome clusters by computing their Mash[55] distance to extant cluster representatives. Comparative metagenomic datasets for wild mouse[33], lab mouse[11], human type I diabetes[56], healthy humans[57], and ocean samples[50] were retrieved from the SRA (accession IDs in S1 Table) and converted to fastq with NCBI’s fastq-dump utility. Metagenomic datasets from wild-type and DNR mice previously studied at the Gladstone Institutes[14] can be found under BioProject PRJNA397886. We used a paired Wilcoxon-rank test to evaluate the change in total hash recruitment by sourmash for the three pairs of reference database settings (default vs SAG-only, default vs combination, combination vs SAG-only). We also tested the difference in the number of species that were assigned more than 5 hashes, as an approximation for species prevalence. For MIDAS, we evaluated differences in median and mean coverage of marker genes, as well as the species prevalence, using the unpaired Wilcoxon-rank test.

## Data Availability

We submitted sequencing runs for 697 SAGs to SRA under BioProject PRJNA481120. Genome assemblies and feature annotations are available in a figshare repository (DOI: 10.6084/m9.figshare.c.4454150)

## Acknowledgments

We thank the staff of the Bigelow Laboratory Single Cell Genomics Center for the generation of single cell genomics data

## Author contributions

F.W. and S.S. performed the mouse work and biological sample extraction. R.S. oversaw SAG generation and sequencing. S.L. performed the computational analyses and wrote the initial draft of the manuscript. K.S.P and R.S. advised and proposed extensions to the analyses. K.S.P. initiated the study. All authors read and approved the final manuscript.

## Captions for Supporting Information

**S1 Fig. Assembly quality improvement with high coverage sequencing**. Multiple metrics are improved when comparing high coverage versus low coverage single cell sequencing data. Facets show the individual metrics assessed: assembly completeness as determined by CheckM, total length of the genome assembly, maximum contig length, total number of reads generated. Numbers over each boxplot represent p-values of paired Mann-Whitney tests.

**S2 Table. Accessions used for taxonomic classifier performance evaluation**. Public data retrieved from SRA and ENA to test the performance of metagenomic classifiers with custom reference databases.

**S3 Table. Results of nonparametric comparisons of taxonomic classifier performance with varying reference databases**. Results of Mann-Whitney tests comparing metagenomic read recruitment metrics for every combination of reference type (default, single-cell genomes only, combined) and test dataset. Two sheets are present in the file, reflecting the results from two different taxonomic classifiers (sourmash and MIDAS)

**S4 Fig. Distributions of taxonomic classifier performance metrics when using the taxonomic classifier sourmash and varying reference databases**. Ridgeline plots representing distributions of 2 metagenomic classifier performance metrics when using sourmash - total number of kmer hashes assigned and number of species with more than 5 hashes (an approximation for prevalence). The plots are faceted by dataset, and each line within the facet reflects one of the three reference database options - default set of genomes available in GTDB release 86, a custom database with single-cell genomes only, and a combined database with the GTDB v86 and single-cell genomes.

**S5 Fig. Distributions of taxonomic classifier performance metrics when using the taxonomic classifier MIDAS and varying reference databases.** Ridgeline plots representing distributions of 3 metagenomic classifier performance metrics when using MIDAS - mean coverage of 15 phylogenetically informative marker genes, median coverage of the same genes, and prevalence (number of samples a species is present in). The plots are faceted by dataset, and each line within the facet reflects one of the three reference database options - default MIDAS v1.2 database, a custom database with single-cell genomes only, and a combined database with the MIDAS v1.2 and single cell genomes.

